# Decoupling Bulk Stiffness and Pore Size in 3D Rat Tail Collagen I Matrices

**DOI:** 10.1101/2025.10.02.680089

**Authors:** Theadora Vessella, Qi Wen, Hong Susan Zhou

## Abstract

The interplay between extracellular matrix (ECM) mechanics and the tumor microenvironment is increasingly recognized as a key driver of cancer progression. Three-dimensional (3D) collagen I culture systems provide more physiologically relevant environments than traditional 2D cultures, yet seperately controlling the structural and mechanical features of 3D matrices remains challenging due to their inherent interdependence. Here, we present a collagen-based approach to decouple bulk stiffness and pore size by varying collagen concentration and polymerization temperature. Collagen concentration was varied to modulate bulk storage modulus, while polymerization temperature was adjusted to alter pore dimensons at comparable collagen concentrations. Using this approach, we generated 3D Collagen I matrices with bulk storage moduli of approximately 80, 228, and 360 Pa while maintaining a similar median pore sizes near 2.5 µm. Conversely, 1.5 mg/mL collagen, median pore size varied from approximately 2.5 to 3.1 µm while bulk storage modulus ramined near 80 Pa, whereas at 3.5 mg/mL collagen, pore size varied from approximately 2.0 to 2.4 µm while bulk storage modulus remained near 350 Pa. We further evaluated the responses of MCF-10A epithelial cells and MDA-MB-231 metastatic breast cancer cells to matrices spanning these sturactual and mechanical conditions. Although the changes in pore size within each stiffness range were modest, they were associated with measurable and statistical differences in cell morphology. These findings demonstrate that even modest changes in collagen network architecture can produce measurable differences in cellular behvarior under comparable bulk mechanical conditions, highlighitng the importance of considering matrix structure in addition to stiffness when investigating cell-matrix interactions.

## 1. Introduction

The extracellular matrix (ECM) plays a central role in both development and disease processes. Through surface receptors, cells sense and respond to the ECM’s chemical composition and physical properties. Dynamic changes in the ECM’s structure and composition can alter its chemical and physical cues, influencing key cellular behaviours such as migration, alignment, proliferation, and morphology. In two-dimensional (2D) continuous substrates, environmental factors such as stiffness, size, density, and the spatial distribution of adhesion sites can be precisely controlled to study their effects on cell migration. However, these properties become more complex and less predictable in three-dimensional (3D) porous hydrogels (1). The most influential parameters affecting cancer cell behaviour in 3D environments include matrix stiffness and confinement, which is often characterized by pore size.

Recent research has shown that changes in stiffness not only accompany disease progression but may actively contribute to it (2). Increased ECM stiffness is a known regulator of tumour proliferation, immune cell infiltration, epithelial to mesenchymal transition and drug delivery (3, 4). Cells exhibit different morphologies, modes of migration, proliferation rates, and nascent ECM secretion in 3D environments when compared with 2D substrates (5, 6).

Similarly, ECM pore size has been shown to be an important factor in determining cell migration patterns, cell phenotype, cell behaviour and protein secretion (7). For instance, when pores are small, confinement can restrict cell locomotion, nutrient diffusion, and removal of waste metabolites. Conversely, when pores are sufficiently large relative to nuclear dimensions, physical confinement is reduced and the cell will undergo invasion (8). However, excessively large pores may also reduce the available surface area for cell attachment (9).

Recent studies in 3D matrices have demonstrated that increased matrix stiffness can enhance cell spreading, tumour outgrowth, ligand densities, intracellular stiffness, and motor activity (10–13). Other works have shown that decreasing pore size, thereby increasing confinement, can affect proliferation, viability, cell morphology and multicellular cluster formation (14–17). However, in many 3D matrix systems, changes in matrix stiffness and pore size remain coupled, making it difficult to delineate whether observed cellular behaviours are driven primarily by stiffness, pore size, or a combination of both. Consequently, cellular responses attributed to an individual matrix property may be influenced by concurrent change in other physical characteristics of the matrix. Developing tuneable 3D matrices that allow physical parameters to be varied over partially overlapping ranges would therefore provide an opportunity to better distinguish the effects of matrix mechanics and architecture on cellular behaviours and disease-relevant phenotypes.

Ideally, cellular processes such as migration, morphology, and matrix remodelling can be studied in engineered cell culture systems in which stiffness changes, and pore size changes can be controlled, independent of one another or over partially overlapping ranges. Recent strategies to modulate 3-D matrix properties have included modifying matrices with alginate (18), using synthetic polymers with tuneable cross-linking densities (19–22), non-enzymatic glycation of collagen (10, 23) and creating inter-penetrating networks comprised of natural proteins (24–27) and other hydrogel materials, like hyaluronic acid (28). These approaches provide valuable insight on generating 3D matrices with tuneable mechanical and structural properties. However, incorporation of additional polymers or chemical modifications can introduce changes in matrix composition, ligand presentation, fibre organization, degradation, or cell-matrix interactions that may contribute to resulting cellular responses. Thus, collagen-based approaches that vary physical properties without introducing additional structural polymers or chemical crosslinkers provide complementary methods for investigating how matrix architecture and bulk mechanics are associated with cell behaviour.

The dimensions and organization of collagen fibrils can be modulated by varying collagen concentration and polymerization temperature under conditions compatible with cell culture. Previous studies have demonstrated that increasing collagen concentration simultaneously increases bulk stiffness, measured as storage modulus (G′), and decreases average pore size (29, 30). Additionally, other reports have demonstrated that decreasing the collagen polymerization temperature to approximately 22°C results in increased pore size (31, 32). This effect is consistent with the temperature dependence of collagen fibrillogenesis, in which polymerization temperature can influence fibril formation, fibre diameter, and network organization. Therefore, polymerization temperature is not expected to selectively alter pore size but rather provides a means of modifying the resulting collagen network architecture. We hypothesized that combining polymerization temperature with collagen concentration could weaken the inherent coupling between pore size and bulk stiffness and enable these properties to be varied over partially overlapping ranges without introducing additional chemical crosslinkers or structural polymers.

Herein, we utilize 3D collagen I matrices polymerized at either low temperature (room temperature, approximately 22°C) or 37°C while simultaneously varying bulk collagen concentration (1.5, 2.5, and 3.5 mg/mL). We characterize 3D collagen I matrix architecture using confocal reflectance microscopy (CRM) and quantify pore size using a three-dimensional Euclidean distance transform (EDT)-based analysis. In parallel, we perform rheological characterization to determine the bulk storage modulus (G′) of the collagen matrices. Together, these measurements allow us to evaluate the relationship between collagen network architecture and bulk mechanical properties across polymerization conditions.

## 2. Results

### 2.1 Temperature- and Concentration-Dependent Collagen Assembly

The lateral growth of collagen fibrils is known to be temperature dependent, and both collagen concentration and polymerization conditions influence the resulting network architecture. Here, we leveraged these properties to modulate pore size and stiffness in collagen matrices without the use of chemical crosslinkers. Neutralized collagen solutions were polymerized in microwells (200 µL volume) either at 37°C (referred to as “37°C”) for 60 minutes or at room temperature for 30 minutes followed by an additional 60 minutes of polymerization at 37°C (referred to as “Room Temp”) (Figure 1A). Confocal reflectance microscopy (CRM) was used to quantify the collagen network architecture without additional sample preparation or drying, which can perturb network structure, thereby enabling assessment of the matrices in their hydrated state (33). Representative CRM images in Figure 1B show the microarchitecture of ColI matrices as a function of the bulk collagen concentration and polymerization temperature. Representative CRM images in Figure 1B show differences in collagen network architecture across matrices containing 1.5, 2.5, and 3.5 mg/mL ColI polymerized at either 37°C or room temperature. Matrices polymerized at 37°C exhibited shorter, more closely spaced collagen fibrils and a visually denser network compared with matrices polymerized at room temperature. At room temperature, the collagen network appeared more open, with larger regions of apparent pore space. Increasing collagen concentration generally resulted in a denser collagen network; however, the 3.5 mg/mL matrix polymerized at room temperature exhibited a comparatively open network with larger visible regions of vacancy.

**Figure 1.**
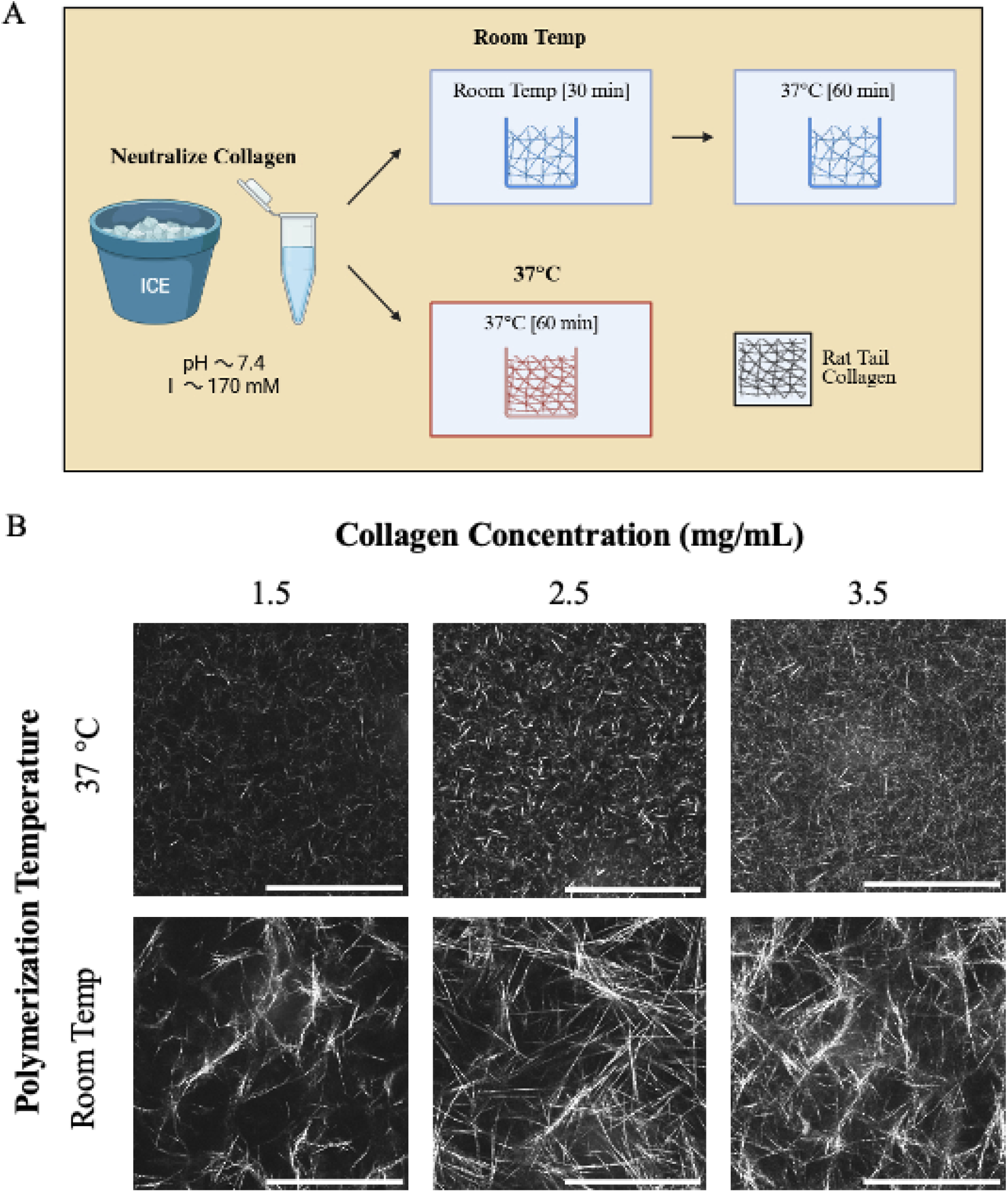
Fabrication and confocal reflectance microscopy of collagen hydrogels. (A) Schematic representatio of the 3D collagen neutralization and gelation process. (B) Maximum projection images of collagen hydrogels captured using confocal reflectance microscopy (CRM) at varying temperatures and concentrations. Experiments were performed in three independent experiments. Scale bars represent 50 μm.

### 2.2 Topological Characterization of 3D ColI Matrices

A 3D image segmentation analysis was applied as described in experimental methods. Briefly, CRM image stack (50 µm height, 0.2 µm step size) were denoised and thresholded prior to application of a 3D Euclidean Distance Transform. Representative binary-segmented CRM images and 3D Euclidean Distance Transform (EDT) images are shown in Figure 2. The CRM image stacks were segmented to distinguish collagen fibrils from pore regions, followed by application of the 3D EDT. The resulting Euclidean Distance Map (EDM) represents the shortest distance from each non-fibril voxel to the surrounding collagen fibrils. Local maxima within the EDM correspond to the radii of the largest sphere that would fit within each respective 3D pore of the collagen scaffold (Figure 2B). Pore diameters were calculated as twice the corresponding radii (2 x radii), and the median of the resulting pore diameter distribution was used as the representative pore size for each collagen matrix. The distribution of pore size diameters was also evaluated to characterize the range and variability of pore dimensions within each matrix condition (Figure 3A). While the median pore diameter reflects the central tendency, histograms reveal the full range of pore sizes, highlighting variability as well as the presence of relatively large or small pores within each matrix condition (Supplemental Figure 1). Three-dimensional pore-size analysis demonstrated that Room Temp ColI matrices exhibited larger median pore diameters compared with 37°C ColI matrices across the collagen concentrations examined (Figure 3A). Room Temp matrices also exhibited visibly thicker fibrillar features compared with 37°C matrices (Figure 3B). Together, these observations indicate that polymerization temperature influences collagen network architecture, including pore dimensions and fibrillar features, rather than selectively altering pore size. Pixel intensity profiles were also used to quantify the spatial distribution of collagen fibrils across the CRM images, providing a qualitative comparison of fibrillar organization between conditions (Figure 3C, Supplemental Figure 2).

**Figure 2.**
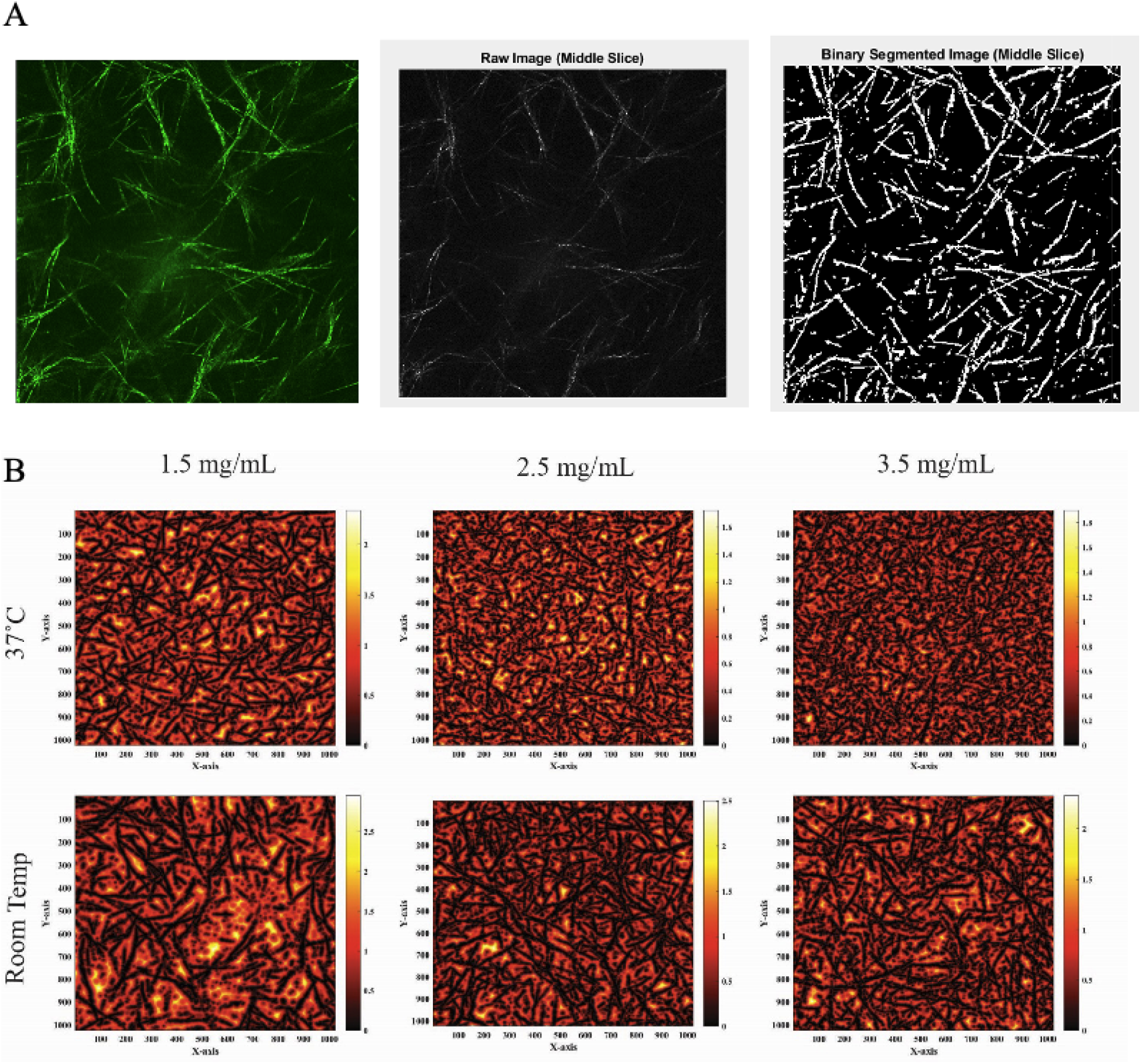
Image processing and Euclidean Distance Transform (EDT) analysis for quantitative determination of ColI matrix pore size. (A) Representative CRM average projection, corresponding raw image, and binar segmented image used for quantitative pore-size analysis of 3D ColI matrices. (B) Representative 3D Euclidea Distance Transform (EDT) images illustrating pore-size analysis across ColI matrices prepared at 1.5, 2.5, and 3. mg/mL collagen concentrations and polymerized at either room temperature or 37°C. Black represents segmente collagen fibres and color gradient represents distance away from collage fibril. Heat map gradient represents EDT detected maxima’s in microns [µm].

**Figure 3.**
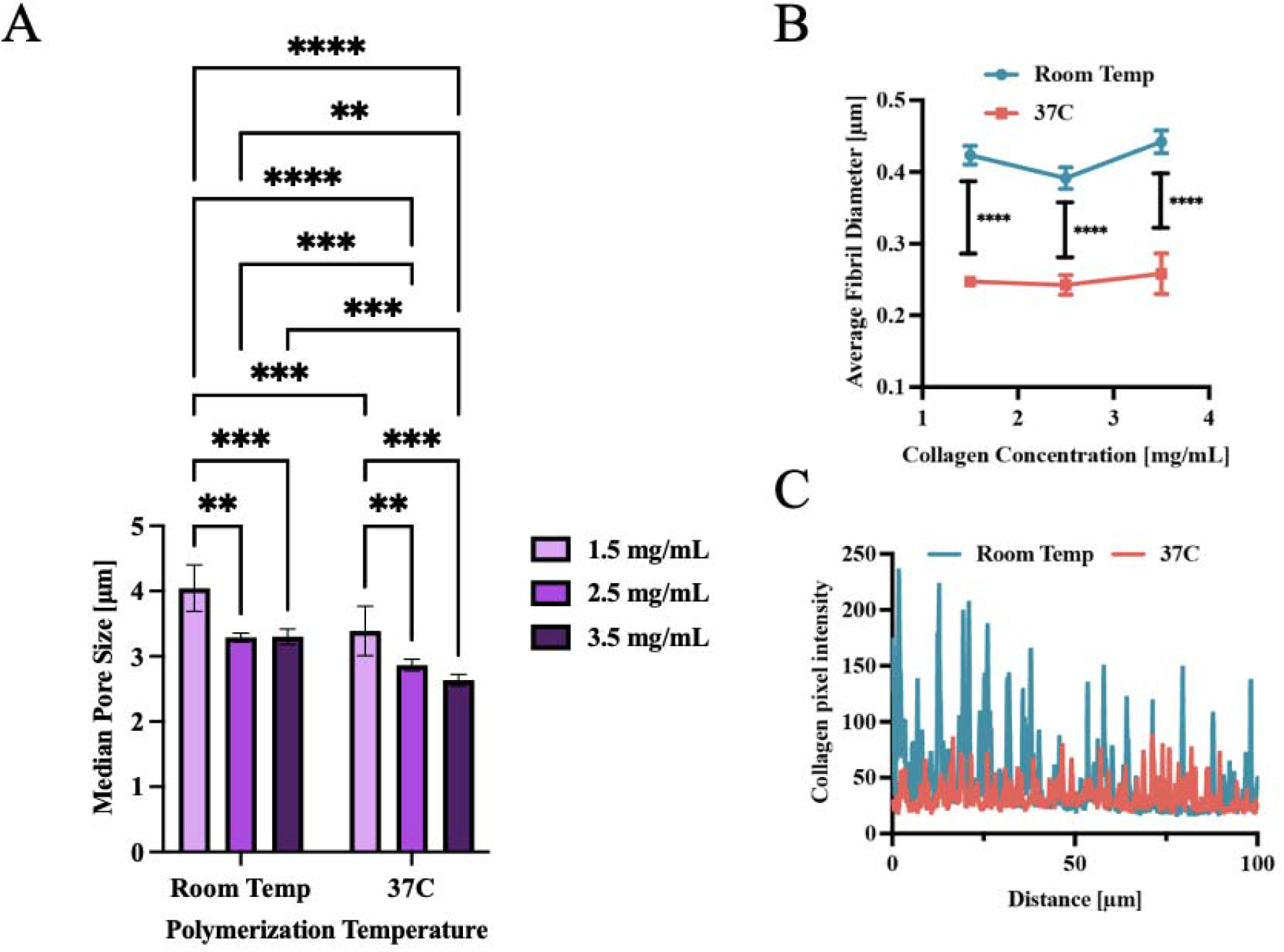
Room temperature collagen polymerization increases median pore size in 3D collagen hydrogels. (A) Comparison of average median pore diameters in each gel under varying collagen concentration and polymerizatio temperature. Experiments were performed in three independent experiments. Two-Way ANOVA followed by Tukey’s post-hoc test, not significant (ns) *P* > 0.05, \**P* ≤ 0.05, \*\**P* ≤ 0.01, \*\*\**P* ≤ 0.001, \*\*\*\**P* ≤ 0.0001. Error bars represent SEM. (B) Line graph represents the mean fibril diameter based on un-paire t-test, not significant (ns) *P* > 0.05, \**P* ≤ 0.05, \*\**P* ≤ 0.01, \*\*\**P* ≤ 0.001, \*\*\*\**P* ≤ 0.0001. Experiments were performed in three independent experiments. Error bars represent SD. (C) CRM image analysis showing the cross-sectional distribution of collagen fibers from 1.5 mg/mL collagen. Additional cross-sectional distributions (2. and 3.5 mg/mL) can be found in Supplemental Figure 2.

### 2.3 Experimental Analysis of ColI Hydrogel Mechanical Properties

We investigated how collagen concentration and polymerization temperature influenced the bulk mechanical properties of the 3D ColI matrices. Consistent with previous studies of 3D collagen matrices, we measured the storage modulus (G′) of the structurally distinct gels using bulk rheology (34–36). To minimize potential effects of polymerization conditions and temperature variation during rheological measurements, rheological characterization was performed on pre-polymerized gels. Coverslips were affixed to the rheometer base, and a 25mm parallel plate geometry was used to perform a time sweep at a fixed frequency of 1 Hz and constant strain of 0.1% for 10 minutes at room temperature (Figure 4A). The storage modulus was calculated as the average of the final 50 measurement points.

**Figure 4.**
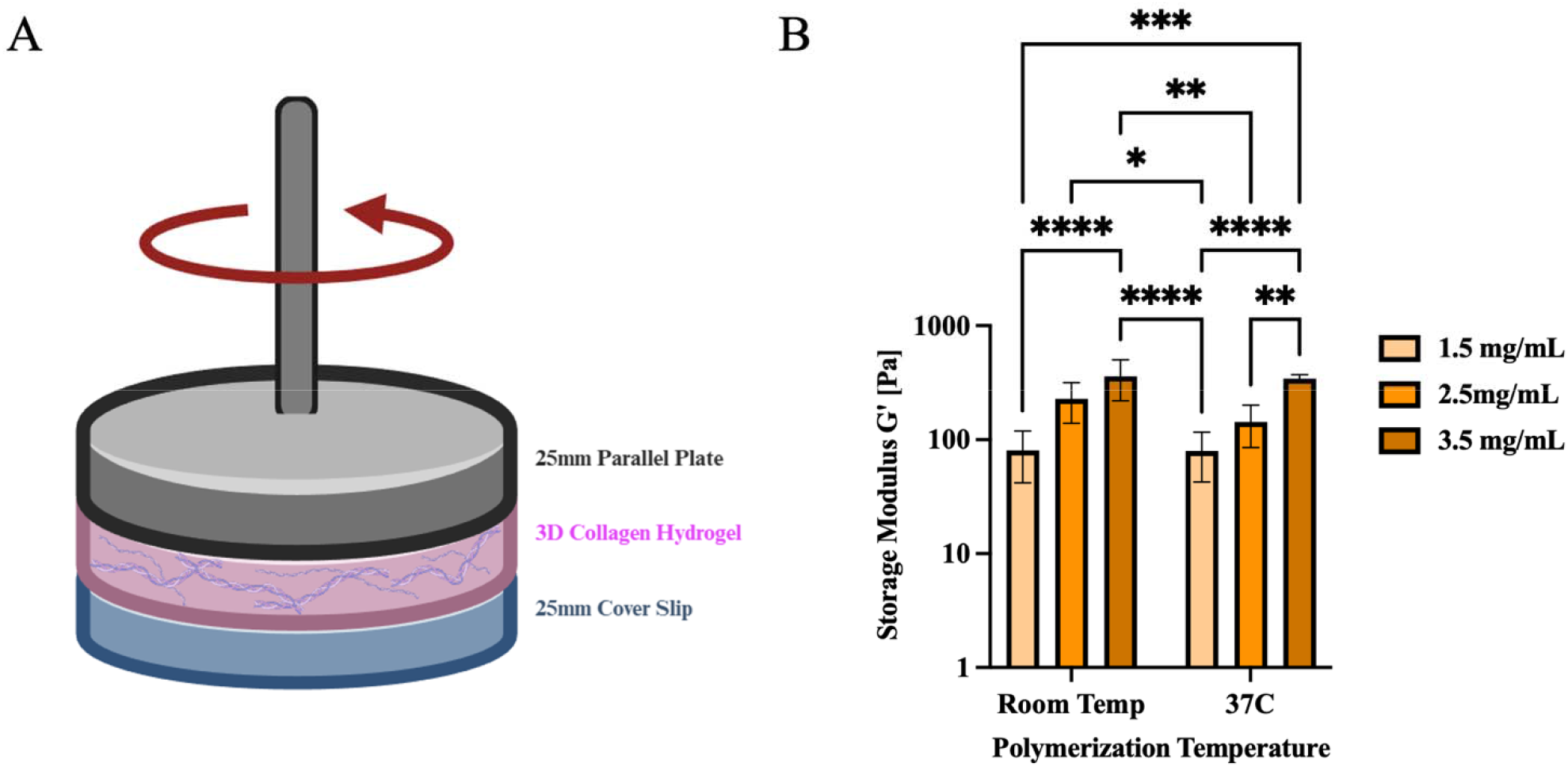
Rheological mechanical characterization of 3D collagen hydrogels. (A) Schematic depiction of rheometer set up for 3D collagen hydrogels utilizing 25mm parallel plate geometry. (B) Mean storage moduli for 37°C and Room Temp collagen hydrogels obtained via rheological measurements (N= 4 - 6 collagen gels). Two-Way ANOVA followed by Tukey’s post-hoc test, not significant (ns) *P* > 0.05, \**p* ≤ 0.05, \*\**p* ≤ 0.01, \*\*\**p*≤ 0.001, \*\*\*\**p* ≤ 0.0001. Error bars represent SD.

When collagen concentration was held constant, we observed no statistically significant difference in G′ between polymerization temperatures (Figure 4B). In contrast, increasing collagen concentration from 1.5 to 2.5 and 3.5 mg/mL resulted in a marked increase in G′. At 37°C, G′ increased from approximately 80 to 230 and 360 Pa, and at room temperature, G′ increased from approximately 80 to 140 and 340 Pa, respectively (Figure 4B). These results indicate that bulk storage modulus was primarily governed by collagen concentration, whereas polymerization temperature had a comparatively smaller effect on the bulk mechanical properties under the conditions examined. Overall, these data demonstrate that collagen concentration provides the primary means of modulating bulk stiffness in these 3D ColI matrices, while changes in polymerization temperature can alter matrix architecture without producing a comparable change in bulk storage modulus.

### 2.4 Decoupling pore size and bulk stiffness via temperature and collagen concentration

Decoupling of pore size and bulk stiffness in 3D ColI matrices was achieved by varying collagen concentration and polymerization temperature. These two parameters altered different aspects of the resulting matrix properties, allowing bulk stiffness and pore size to be varied over partially overlapping ranges. Bulk stiffness was primarily modulated by increasing collagen concentration (e.g., from 1.5 to 2.5 mg/mL), which increased collagen content and bulk storage modulus, whereas decreasing the polymerization temperature from 37°C to room temperature increased pore size. This approach enabled modulation of bulk stiffness while maintaining comparable median pore sizes across selected conditions (Table 1).

**Table 1.** Independent Tuning of Stiffness in 3D Collagen Hydrogels.

| Tune Stiffness |  |  |  |
| --- | --- | --- | --- |
| Collagen Concentration [mg/mL] | Polymerization Temperature | Storage Modulus [Pa] ± Std Dev | Median Pore Size [$\mu\text{m}$] ± Std Dev |
| 1.5 | 37°C | 80 ± 37 | 2.5 ± 0.23 |
| 2.5 | Room Temp | 228 ± 90 | 2.5 ± 0.13 |
| 3.5 | Room Temp | 360 ± 140 | 2.4 ± 0.09 |

Using this approach, we generated three ColI matrices with comparable median pore sizes (∼2.5 μm) but varying bulk storage moduli of 80, 228, and 360 Pa (Table 1, Figure 5A, Supplemental Figure 3). Conversely, at a fixed collagen concentration, polymerization temperature was varied to alter pore size while maintaining a comparable bulk storage modulus (Table 2, Supplemental Figure ). Specifically, at low collagen concentration (1.5 mg/mL), median pore size varied from 2.5 to 3.1 μm while bulk storage modulus remained near 80 Pa. At high collagen concentration (3.5 mg/mL), median pore size varied from 2.0 to 2.4 μm while bulk storage modulus remained near 350 Pa (Table 2, Figure 5B-C). Together, these results demonstrate substantial decoupling of bulk stiffness and pore size within defined ranges of 3D ColI matrix conditions without relying on chemical crosslinkers or synthetic polymer additives.

**Figure 5.**
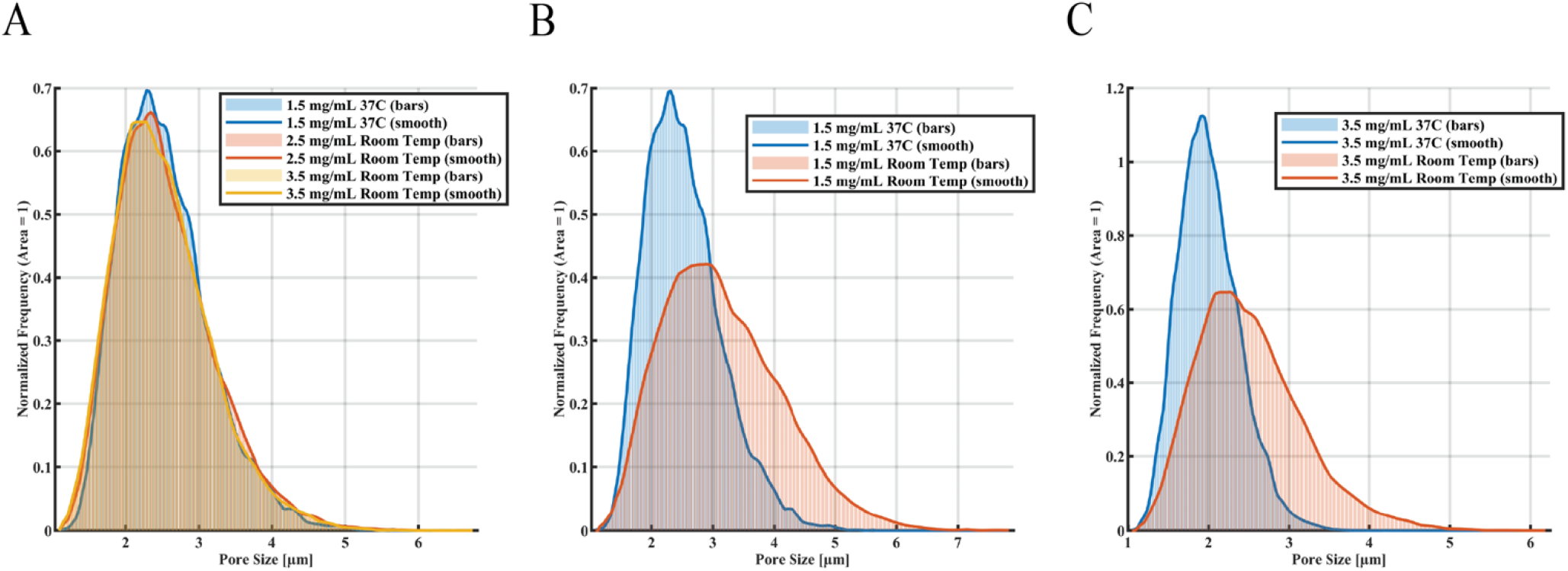
Independent tuned pore size and stiffness in 3D ColI matrices. Average normalized pore size distributions of (A) independently tuned stiffness with no change to pore distribution. Independently tuned pore size at both (B) low- (1.5 mg/mL) and (C) high- (3.5 mg/mL) collagen concentration.

**Table 2.**
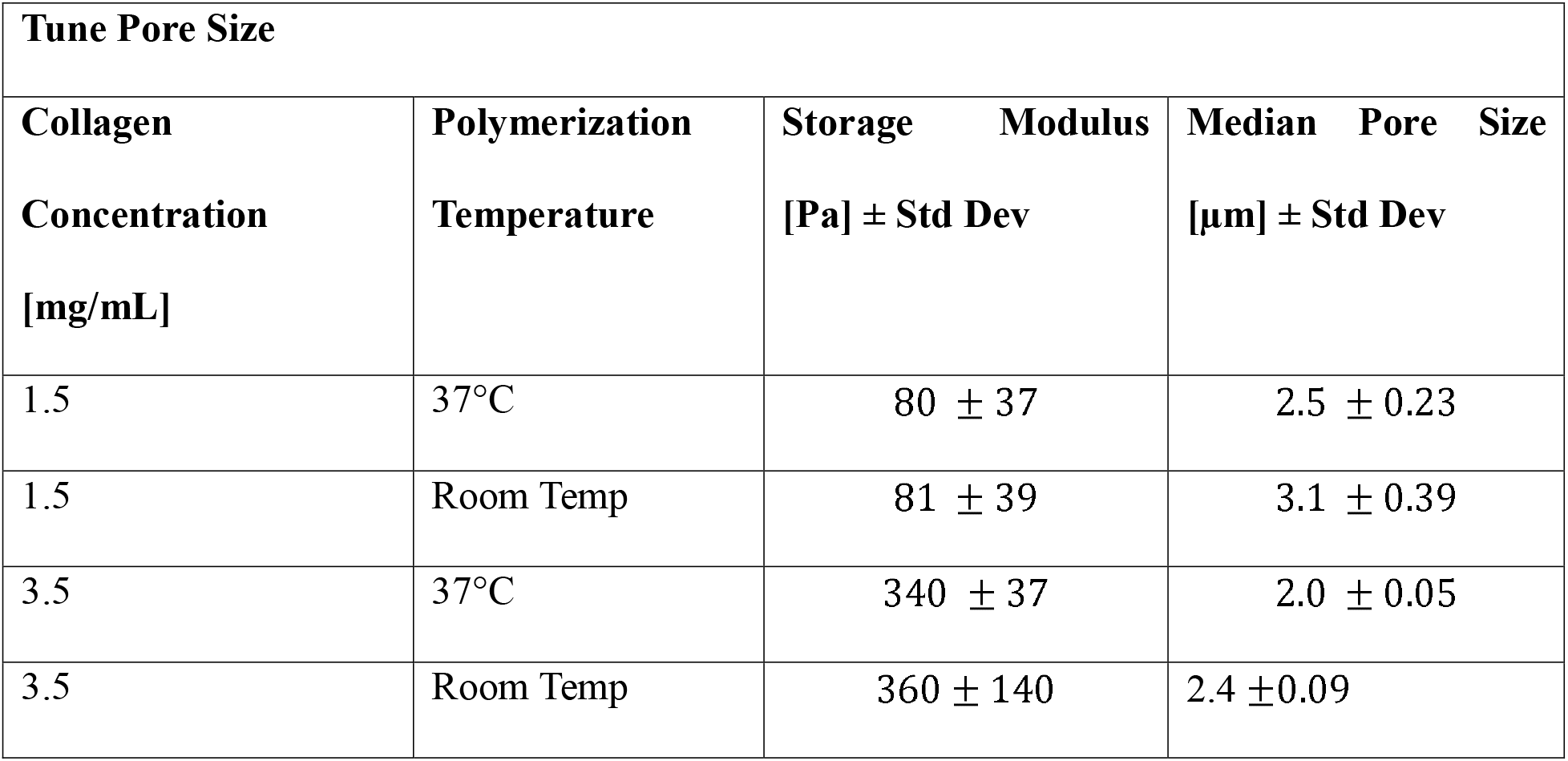
Independent Tuning of Pore Size in 3D Collagen Hydrogels.

### 2.5 Impact of ColI stiffness and pore size on cell morphology and cluster formation

To investigate how extracellular matrix (ECM) properties influence cell behavior, we evaluated cell responses within 3D ColI matrices with defined bulk mechanical and structural conditions. Based on the matrix characterization described above, we evaluated the effects of matrix stiffness using ColI matrices with comparable median pore sizes (∼2.5 µm) and varying bulk storage moduli of 80, 228, and 360 Pa (Table 1). To evaluate the effect of pore size, we compared gels of similar stiffness but different pore sizes: 80 Pa gels with median pore sizes of 2.5 and 3.1 um, 350 Pa gels with median pore sizes of 2.0 and 2.4 µm.

Two breast cell lines with distinct migration characteristics and phenotypes were selected: MDA-MB-231 cells exhibit mesenchymal-like migration, whereas the non-tumorigenic epithelial cell line MCF-10A exhibits collective migration. These cell lines are commonly used models for investigating biological and biophysical responses to the extracellular environment. Single cells were embedded within the ColI matrix, polymerized, and cultured for 4 days. In acellular controls, collagen pore size remained stable over four days, with no detectable changes observed (Supplemental Figure 4). After 4 days, samples were fixed and stained with Alexa Fluor 488 phalloidin, and confocal microscopy was used to characterize cell cluster formation and individual cell morphology (Figure 6). MCF-10A cells formed larger clusters than MDA-MB-231 cells (Figure 6A). MCF-10A cluster formation decreased with increasing matrix stiffness (Figure 6B). In contrast, MDA-MB-231 cells remained predominantly as individual cells across the stiffness conditions examined.

**Figure 6.**
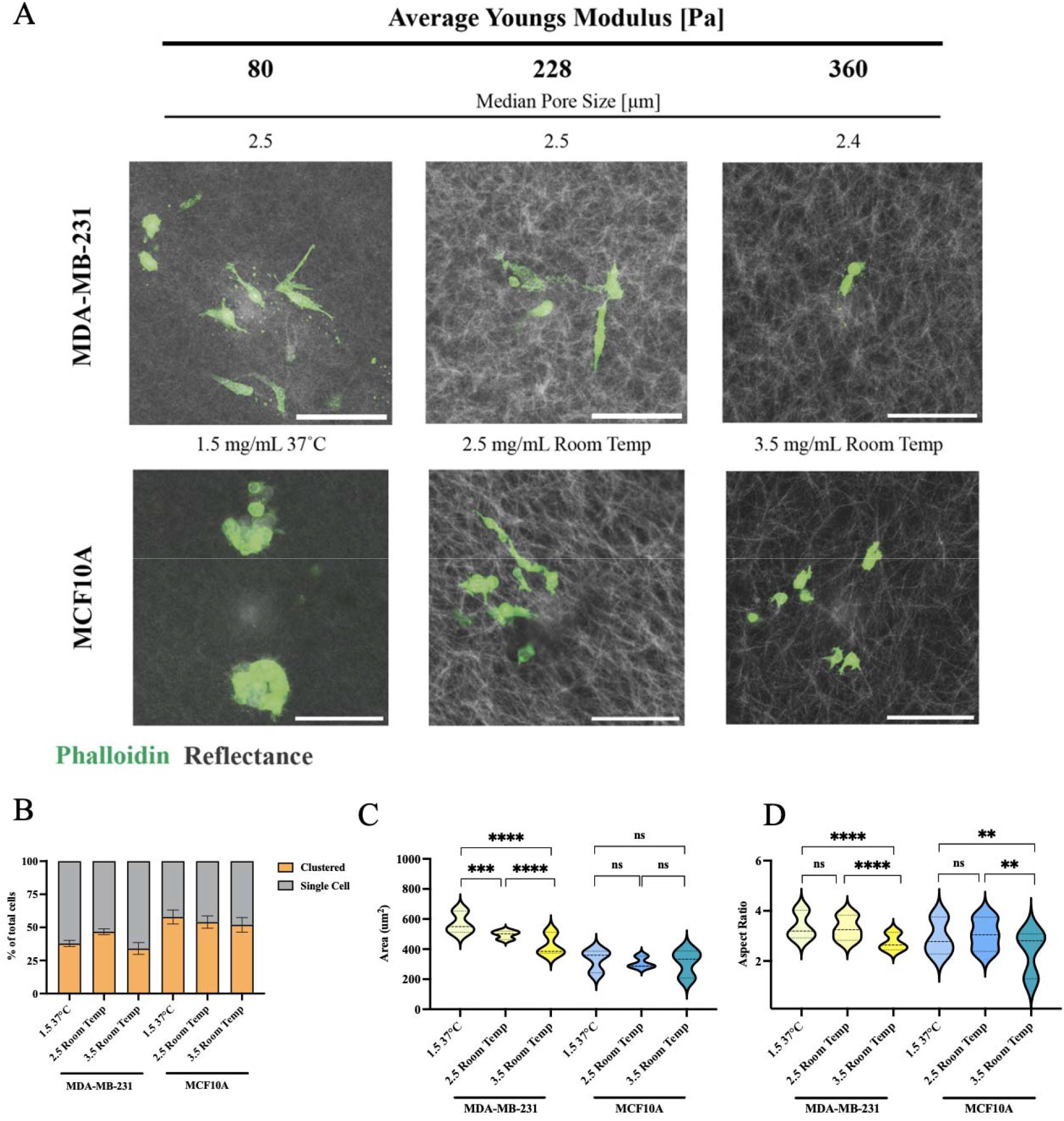
Impact of ColI matrix stiffness on cell morphology and cluster formation. (A) CRM images of MDA-MB-231 breast cancer cells and MCF-10A breast epithelial cells cultivated within 3D ColI matrices with stiffnesses of 80, 228, and 360 Pa for 4 days. Scale bars represent 100 µm. (B) Quantitative analysis of cluster formation of MDA-MB-231 cells and MCF-10A cells, whereby cells without contact to other cells were counted as single cells (N = 3 independent gels per condition; MDA-MB-231 *n* = 455–1125 cells, MCF-10A *n* = 387–687 cells). One-way ANOVA followed by Tukey’s post-hoc test, not significant (ns) *p*> 0.05, *p* ≤ 0.05, \**p*≤ 0.01, \*\**p* ≤ 0.001, \*\*\**p* ≤ 0.0001. Error bars represent SEM. Quantitative analysis of MDA-MB-231 breast cancer and MCF-10A breast epithelial cell area (C) and aspect ratio (D) as a function of stiffness (N = 3 independent gels per condition; MDA-MB-231 *n* = 455–1125 cells, MCF-10A *n* = 387–687 cells). Significance was tested between samples of varying stiffness and similar pore size. Nested one-way ANOVA followed by Sidak’s multiple-comparisons test was used, with individual measurements nested within biological gels. Not significant (ns) *p*> 0.05, *p* ≤ 0.05, \**p*≤ 0.01, \*\**p* ≤ 0.001, \*\*\**p* ≤ 0.0001. Error bars represent SEM.

Analysis of individual cell morphology revealed distinct responses to matrix stiffness between the two cell types. MDA-MB-231 cells exhibited decreased cell area and aspect ratio with increasing gel stiffness (Figure 6C–D). In contrast, MCF-10A cells maintained smaller cell areas and aspect ratios than MDA-MB-231 cells across all stiffness conditions. Although MCF-10A cell area remained largely unchanged, aspect ratio decreased and circularity increased with increasing matrix stiffness (Figure 6C–D, Supplemental Figure 5).

The effect of pore size on cellular behavior was also examined under both low (1.5 mg/mL) and high (3.5 mg/mL) collagen concentrations (Figure 7A). Across all conditions, MCF-10A cells consistently formed larger clusters than MDA-MB-231 cells (Figure 7B, E). At low collagen concentration (1.5 mg/mL, ∼80 Pa), MDA-MB-231 cells exhibited decreased area and aspect ratio and increased circularity with decreasing pore size from 3.1 µm to 2.5 µm (Figure 7F–G, Supplemental Figure 5). In contrast, MCF-10A cells exhibited increased area and decreased aspect ratio with decreasing pore size. At high collagen concentration (3.5 mg/mL, ∼350 Pa), MDA-MB-231 cells exhibited increased area and aspect ratio with decreasing pore size from 2.4 µm to 2.0 µm, while MCF-10A cells displayed the opposite trend, with cell area decreasing with decreasing pore size over the same range.

**Figure 7.**
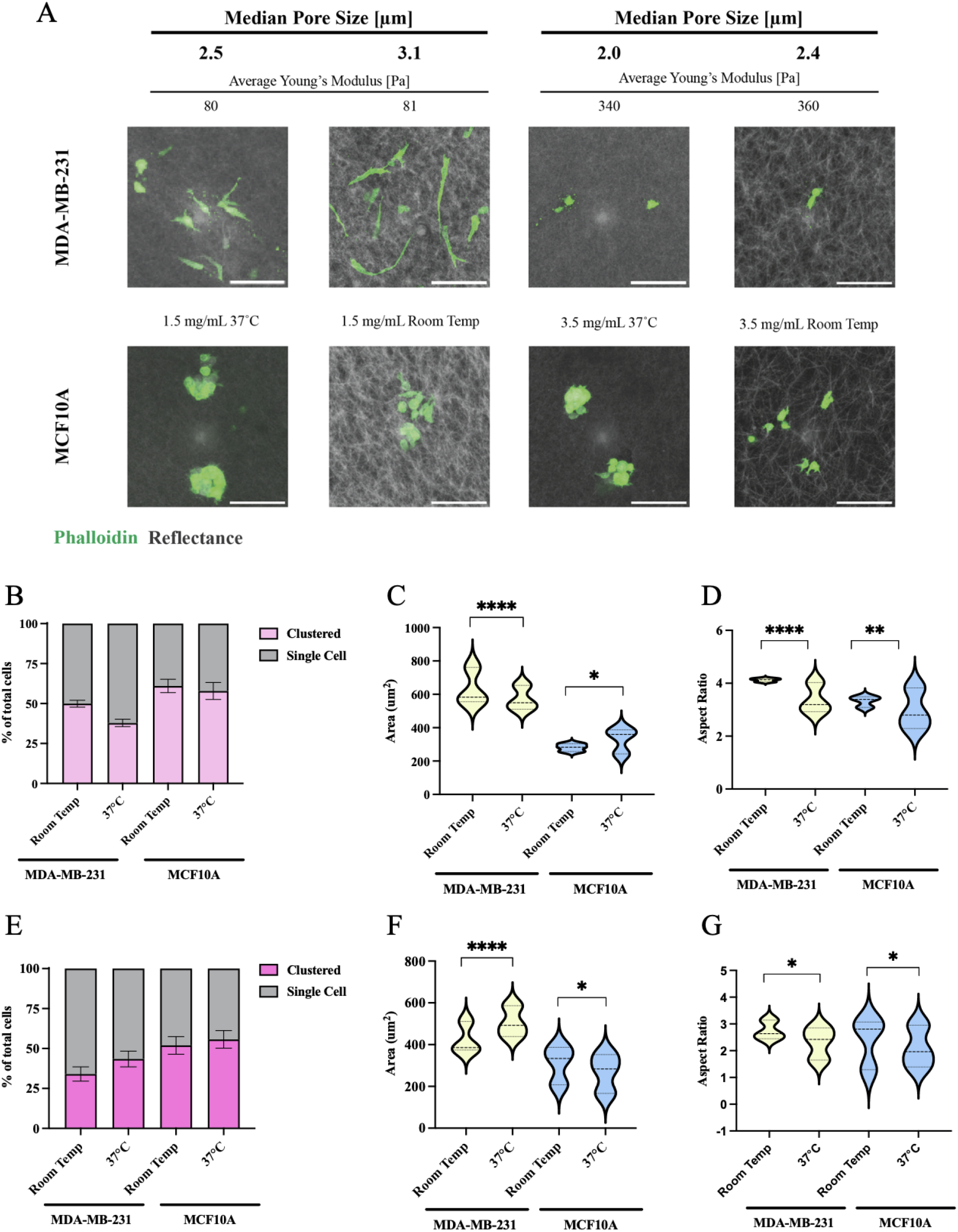
Impact of ColI pore size on cell morphology and cluster formation. (A) CRM images of MDA-MB-231 breast cancer cells and MCF-10A breast epithelial cells (green) cultivated within 3D ColI matrices (grey) of both low collagen concentration (1.5 mg/mL) and varying median pore size from 2.5 µm to 3.1 µm and high collagen concentration (3.5 mg/mL) with varying median pore size from 2.0 µm to 2.4 µm for 4 days. Scale bars represent 100 µm. Quantitative analysis of cluster formation of MDA-MB-231 cells and MCF-10A cells at both low collagen concentration (B) and high collagen concentration (E). Cells without contact to other cells were counted as single cells (grey) (N = 3 independent gels per condition, with 5–15 z-stacks acquired per gel. Quantitative analysis of MDA-MB-231 breast cancer and MCF-10A breast epithelial cell area and aspect ratio as a function of pore size for both low collagen concentration (C–D) and high collagen concentration (F–G) (N = 3 independent gels per condition; MDA-MB-231 *n* = 455–1125 cells, MCF-10A *n* = 387–687 cells). Significance was tested between samples of varying pore size and similar stiffness. Nested one-way ANOVA followed by Sidak’s multiple-comparisons test was used, with individual measurements nested within biological gels. Not significant (ns) *p* > 0.05, *p* ≤ 0.05, \**p* ≤ 0.01, \*\**p* ≤ 0.001, \*\*\**p* ≤ 0.0001. Error bars represent SEM.

## 3. Discussion

Cancer cell invasion is strongly regulated by the biophysical properties of the ECM, including its microstructure and mechanical stiffness. However, these properties are often interdependent, making it difficult to recapitulate the native ECM architecture and independently examine their effects on cell behavior. This problem is further compounded in common approaches for tuning the mechanics of 3D collagen matrices, which often rely on chemical-crosslinkers. The current study demonstrates a method to substantially decouple matrix stiffness and pore size without the need for chemical crosslinkers or polymer additives. Our findings show that bulk stiffness can be tuned from 80 Pa to 360 Pa by varying collagen concentration while maintaining a comparable median pore size of 2.5 μm through polymerization temperature control. Conversely, at a fixed collagen concentration (e.g., 1.5 mg/mL) where bulk stiffness remained near 80 Pa, adjusting the polymerization temperature allows pore size to be modulated from 2.5 μm to 3.1 μm. Cellular studies further demonstrated distinct responses of metastatic and non-tumorigenic breast cancer cells to changes in matrix stiffness and pore size. Together, these results highlight the utility of controlling polymerization temperature and protein concentration to substantially decouple 3D matrix stiffness and pore size within defined ranges, enabling further precision to study cellular responses to mechanical cues in 3D environments.

Methods to alter polymerization temperature for pore size control have been explored previously in the literature. Several reports have described methods to alter fiber diameter, or pore size, of 3D ColI matrices using decreased polymerization temperatures (31, 37). While these studies demonstrate that lower polymerization temperatures can increase pore size, they do not isolate stiffness from pore size without introducing chemical crosslinkers or other polymers into the ColI network. Additionally, many of these approaches simultaneously alter pH and ionic strength outside of physiological conditions, which likely alter collagen electrostatic interactions and further confound mechanical and structural properties. Importantly, maintaining collagen as the sole matrix component minimizes the introduction of additional biochemical or structural cues that may independently influence cellular behavior. While approaches incorporating hyaluronic acid, glycation, or other matrix modifications can alter mechanical properties, these modifications may also change ligand presentation, matrix composition, or fibrillar organization. By avoiding additional matrix components or chemical modifications, the present approach reduces the number of concurrent biochemical and structural variables introduced during matrix tuning, allowing observed cellular responses to be more directly interpreted in the context of bulk stiffness and pore size. It is important to note that the stiffness of cancerous and healthy breast ECM has been reported to range from approximately 200 Pa to 4000 Pa, while our 3D ColI gels range from 80 to 350 Pa (38). Thus, our system spans a substantial range of bulk stiffness within the lower range of reported breast ECM stiffnesses. Within these relatively small variations in matrix stiffness, cells exhibit measurable mechanosensitive responses, suggesting that changes in bulk stiffness within this range can regulate cell behavior and merit further investigation.

We also define additional conditions to maintain stiffness, enabling pore size modulation. At these parameters, we also observe a distribution of pore sizes ranging from 1 μm to 7 μm, with median pore sizes between 2.0 and 3.1 μm under comparable stiffness conditions. Given the inherent heterogeneity and occasional unpredictability of 3D ColI, it is important to note that variation in polymerization temperature may introduce increased variability and that changes in polymerization temperature may affect not only pore size, but also other aspects of collagen architecture, including fibril thickness, fiber organization, and network connectivity. While our approach enables modulation of pore size within comparable bulk stiffness conditions, these other structural parameters may also vary and should be considered when interpreting the effects of matrix architecture. Overall, these results provide a biomimetic 3D microenvironment that is consistent with other reports (39).

While prior studies have primarily relied on indirect metrics such as 2D imaging or porosity to assess the matrix structure, our method directly quantifies 3D pore size, offering greater precision when characterizing the pericellular matrix dimensions. Pore size or mesh size is an important parameter for characterizing the structure of biopolymeric filamentous networks, as it can influence cellular behavior like adhesion, polarization, and migration (8, 40). Due to the randomness of biological networks and the heterogeneity of collagen matrices, the extraction of physical parameters via confocal microscopy is not straightforward and remains a non-trivial task. To overcome this, we applied a 3D Euclidean distance transformation to segmented collagen networks to quantify the largest diameter that can fit within each pore. This method provides an estimate of the spatial constraints experienced by cells as they migrate through the matrix.

Stiffness was substantially decoupled from pore size, and MDA-MB-231 cells responded by decreasing their cell area and aspect ratio with increasing stiffness. Previous studies have shown that these cells migrate more slowly in stiffer matrices, which may indicate that they require more time to deform and remodel the surrounding extracellular matrix (41). Previous studies have reported that MDA-MB-231 cells exerted approximately 10-fold higher traction forces on soft gels compared to stiff ones, a surprising finding given that many cell types typically become more contractile on stiffer substrates. Additionally, it is well established that traction forces tend to decrease with decreased cell spreading area (42–46). This suggests that MDA-MB-231 cells may adopt a different migration or mechanosensing strategy in response to increased stiffness, possibly involving amoeboid migration, which relies more on cortical tension and cell deformability than on traction forces. Amoeboid migration in MDA-MB-231 cells has been associated with a rounder cell morphology. One study investigating MDA-MB-231 cells under varying collagen densities, which also increases stiffness and decreases pore size, reported a reduction in both cell area and migration speed, along with the emergence of more rounded, amoeboid-like cell morphologies. While these findings are consistent with our observed changes in cell morphology, migration mode was not directly assessed in the present study (47). These findings therefore suggest a possible relationship between increased matrix stiffness, reduced cell spreading, and a more rounded morphology, although the underlying migration strategy was not directly evaluated.

Pointcloix et. al. further reported that the roundedness of MDA-MB-231 cell generated tensional forces through 1-integrin adhesion and the absence of lamellipodial extensions (48). Similarly, other studies have shown that an increased matrix stiffness can reduce invadopodia formation thereby limiting metastatic invasion, compared to softer 3D environments (49). Integrin-mediated singling has also been shown to play a critical role in activating downstream pathways, including the Rho/ROCK pathway, which is a known regulator of actin dynamics and cellular contractility. These signaling mechanisms may contribute to the observed cellular behavior, although β1-integrin signaling, Rho/ROCK activation, invadopodia formation, and migration mode were not directly evaluated in the present study. Therefore, these mechanisms should be considered potential explanations supported by previous literature rather than direct conclusions from the current data. Together, these findings suggest that the reduced cell areas observed in MDA-MB-231 cells may contribute to decreased traction forces, which could be consistent with ameboid migration patterns.

MCF10A cells were less likely to form large clusters in stiffer 3D ColI matrices, suggesting that matrix stiffness influences collective cell organization. This behavior contrasts with softer matrices, where a more permissive mechanical environment may support collective behavior, allowing cells to aggregate and form tumor-like structures. A previous study has shown that decreased cluster formation in stiff matrices shifts from non-tumorigenic MCF10A cells led to an increase in invasive sites presenting higher levels of vimentin, a key regulator in mechanosensation (50). However, while no significant change in cell area was observed in MCF10A cells, aspect ratio decreased in stiffer matrices, indicating a shift towards a more rounded morphology.

Cellular interactions with the 3D ColI matrices provide further insight into how biophysical properties, including stiffness and pore size, influence cellular behavior. Cell–matrix interactions encompass multiple processes, including protrusion extension, adhesion formation, and matrix remodeling, all of which can be influenced by the local pore architecture.

Based on the data, increasing the collagen concentration (low vs. high) while varying pore size within the defined stiffness conditions differentially influenced the morphological response of MDA-MB-231 and MCF-10A cells. For MDA-MB-231 cells in low collagen concentration, increasing the pore size led to an increase in aspect ratio and cell area. This suggests that in environments with less collagen density and larger pores size, MDA-MB-231 cells may better extend their protrusions due to reduced steric hindrance, facilitating a more elongated shape, although migration or invasion was not directly assessed in this study. Conversely, in a high collagen concentration, increasing pore size resulted in decreased cell areas and aspect ratios.

One possible explanation is that the larger pores in an otherwise dense collagen matrix create fiber-sparse regions, causing cells to retract and decrease aspect ratios due to insufficient anchoring. In contrast, smaller pores within high concentration gels may provide enhanced fibril alignment and bundling for guidance that supports their elongation. However, differences in fibril thickness, collagen organization, and available adhesion sites may also contribute to these concentration-dependent responses, as these local matrix properties were not directly measured in this study. Additionally, under these 3D conditions, the acquired more rounded morphology at higher collagen concentration may reflect cellular adaptation to higher matrix density (47, 51). These findings highlight the ability of cells to adapt their morphology to differences in collagen concentration and pore size, although the underlying mechanisms require further investigation.

In contrast, the morphological response of MCF-10A cells was also dependent on collagen concentration, with changes in pore size producing different responses under low and high collagen conditions. At low collagen concentration, decreasing pore size increased cell are and aspect ratio, whereas at high collagen concentration, decreasing pore size decreased cell area. The ability of MCF-10A to form large, multicellular clusters appears to be consistent with their ability to proliferation and form spheroid-like clusters. This observation is consistent with other reports, where MCF10A cells proliferate to form spheroid-like clusters (52) after 4 days in culture and that reduced pore size is associated with decreased single-cell spreading. It is also known that MCF10A display stable cadherin junctions and high E-Cadherin expression, regulating strong cell-cell adhesion and reducing the potential of the epithelial to mesenchymal transition (EMT) (53). Similarly to the MDA-MB-231 cells, MCF-10A also exhibited reduced cell area and aspect ratios with increasing collagen content. This suggests that denser matrices impose greater physical constraints on cell spreading and elongation across both malignant and non-tumorigenic cell lines.

While bulk rheological measurements provide a useful measure of the overall mechanical properties of the 3D ColI matrices, they do not fully capture the local mechanical cues experienced by embedded cells. Local fiber stiffness, network connectivity, fibrillar organization, and ligand presentation may also influence cellular mechanosensing and behavior. Additionally, the present study primarily evaluated the storage modulus and did not characterize other mechanical properties, such as loss modulus, nonlinear strain stiffening, or stress relaxation. Therefore, the observed cellular responses should be interpreted in the context of bulk matrix stiffness, while recognizing that local and time-dependent mechanical properties may also contribute to cellular behavior.

Given that the diffraction-limited resolution of optical microscopy may lead to overestimation in the diameter of collagen fibrils smaller than 200 nm, results in Figure 3C should be interpreted qualitatively. A limitation of CRM is that it cannot detect fibers oriented at large angles relative to the imaging plane (54), as the technique relies on reflected light. As a result, CRM-based measurements may overestimate pore size (46). Additionally, although acellular structural stability of the 3D ColI architecture was assessed over 10 days (data not shown), the stability of pore size, fiber organization, and stiffness in the presence of cells was not directly assessed. Cell-mediated matrix remodeling over the 4-day culture period may therefore contribute to the observed cellular responses.

In summary, we provide a strategy to substantially decouple stiffness and pore size in 3D ColI matrices utilizing collagen concentration and polymerization temperature. Additionally, we utilized a 3D EDT analysis to quantify 3D pore size within the 3D ColI matrices, rather than relying on 3D analysis that may overestimate pore size. We show that MDA-MB-231 and MCF-10A cells line respond differently in terms of cluster formation and cellular morphology under varying stiffnesses and pore sizes. It is important to note that these changes may vary between cell lines. As such, this developed method could serve as an important tool to study cellular responses to defined matrix properties in 3D tumor microenvironments and could be used to inform future tissue engineering and personalized medicine screening platforms.

## 4. Conclusion

In this study, we provide a strategy to substantially decouple stiffness and pore size in 3D ColI matrices utilizing collagen concentration and polymerization temperature. Additionally, we utilized a 3D EDT analysis to quantify 3D pore size within the 3D ColI matrices, rather than relying on 2D analysis that may overestimate pore size. These results highlight the importance of controlling mechanical and architectural features to elucidate their distinct effects on cell morphology. By varying collagen concentration and polymerization temperature, we were able to generate 3D ColI matrices with defined combinations of bulk stiffness and pore size without introducing chemical crosslinkers or synthetic polymer additives. This approach enabled us to examine cellular responses under matrix conditions in which stiffness and pore size could be substantially decoupled within defined ranges. MDA-MB-231 and MCF-10A cells exhibited distinct changes in morphology and cluster formation across these matrix conditions, further demonstrating that differences in the mechanical and architectural properties of the 3D ECM can influence cellular behavior. Overall, our results provide a reliable platform for studying defined mechanical and architectural cues in a 3D ECM biomimetic platform. This approach may be useful for investigating how individual and combined properties of the tumor microenvironment contribute to cancer cell behavior and may provide a foundation for developing more physiologically relevant 3D models for tissue engineering and therapeutic screening. Future studies should further investigate the contributions of collagen fiber organization, local matrix mechanics, and cell-mediated matrix remodeling to cellular responses within these 3D environments. Additional characterization of migration, invasion, and mechanosensitive signaling could further clarify how these matrix properties regulate cancer cell behavior.

## 5. Experimental Methods

### 5.1 Preparation of 3D ColI Matrices

3D collagen matrices were formed using rat tail type I collagen (Advanced Biomatrix, RatCol™, 4 mg/mL). Final collagen concentrations of 1.5, 2.5 and 3.5 mg/mL were adjusted by mixing the manufacturer-provided neutralization solution, sterile water and phosphate buffer to achieve constant pH (∼7.4) and constant ionic strength (∼170mM). ColI solutions were prepared and kept on ice (4°C) to prevent polymerization. Subsequently, 200µL of collagen solution was transferred onto imaging or rheological characterization glass slide, as needed. To initiate polymerization, “37°C” gels were placed at 37°C for 1 hour and the “Room Temp” gels were placed at room temperature (∼22°C) for 30 minutes followed by 37°C, 5% CO and 95% humidity for 1 hour. To maintain proper hydration, 1X phosphate buffer or cell culture media was added to gels after 30 minutes of polymerization.

### 5.2 Rheological Preparation of 3D ColI Matrices

Glutaraldehyde-treated glass coverslips were used during rheological testing to minimize slip, Circular 25 mm glass coverslips were first cleaned and silanized by incubating them in a 1% solution of 3-aminopropyl-trimethoxysilane (Alfa Aesar, Haverhill, MA) for 10 minutes. They were then treated with 0.5% glutaraldehyde (Amresco, Solon, OH), rinsed thoroughly with deionized water, and allowed to dry. The collagen gel solution was prepared on ice as described previously. A volume of 500 µL of the collagen solution was applied to each treated coverslip and allowed to polymerize. Rheological testing was performed immediately after polymerization.

### 5.3 Rheological Assessment on 3D ColI Matrices

The bulk mechanical stiffness of collagen gels were measured using an Anton Paar MCR 302 rheometer with a temperature-controlled stage. Gel disks (25 mm in diameter and approximately 1 mm in thickness) were prepared on circular glass coverslips. Measurements were performed using a parallel-plate geometry with a 25 mm serrated upper plate. constant normal force of 0.25 N was applied to ensure consistent contact between the rheometer plate and the gel surface, with the resulting gap size varying between samples based on sample thickness. The storage modulus (G’) measured under oscillatory shear strain ( 0.1% in amplitude and 1 Hz in frequency).

### 5.4 Confocal Reflectance Microscopy for 3D Collagen Architecture Analysis

A confocal laser scanning microscope in reflectance mode was used to obtain images of fibrillar collagen microstructure for quantitative assessment. All samples were prepared in Bio-Labs 16 culture well with removable #1.5 chambered coverglass. Briefly, collagen hydrogels were neutralized and allowed to self-assemble at desired polymerization temperature as previously described. Immediately after polymerization, images were collected in reflectance mode with a Leica Stellaris SP8 using a 63X oil immersion objective (N.A = 1.40). Each z-stack began at least 10 µm above the cover glass and consisted of 50 slices with 0.2 µm spacing between slices (10 µm total thickness). Three stacks were acquired per sample.

### 5.5 Collagen Architecture Quantification

A 3D pore-size analysis method was adapted as described by Fischer at al. (55). Critical metadata from the image stack, such as voxel size, were directly obtained from the Leica microscope. To account for scattering and absorption due to sample heights and large travel lengths of exciting and emitted light, image segmentation of fibril and non-fibril volumes was performed for each image in the stack. Before segmentation, total variation denoising and a Gaussian filter were applied to reduce image noise while preserving collagen fiber edges. Adaptive local thresholding was subsequently applied to segment collagen fibrils from the surrounding pore space, accounting for variations in signal intensity within the image stack. The segmentation resulted in a 3D binary volume where the black area represents pores and white represents collagen fibers. A 3D Euclidean Distance Transform (EDT) was applied to the non-fibril part of the 3D binary volume. The Euclidean Distance Map (EDM) represents the shortest distance of each non-fibril pixel to the neighboring collagen fibrils (Figure 4B). A Gaussian filter was applied to the EDM, and pores along the edge were excluded. Local maxima were determined from the filtered EDM. The minimum detectable pore size is defined by the numerical aperture of the objective lens. The maxima represent the radii of the largest sphere that would fit into each respective 3D pore of the collagen scaffold. The pore diameters were calculated as 2 x radii, and the actual pore size, ζ, is given by the median of all pore diameters of the collagen scaffold sample, as given in Eq. 1

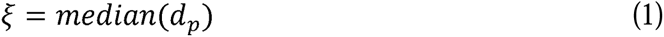

To estimate the fiber diameters, the home-built image processing procedure was utilized as described by Franke et al. (14) equipped with an erosion algorithm and autocorrelation analysis, respectively. Briefly, this method automates the image segmentation into pore and fibril segments for each XY-image of the image stack and evaluates the mean fibril diameter via autocorrelation.

### 5.6 Breast Epithelial and Metastatic Breast Cancer Cell Culture

Human mammary epithelial cell line, MCF10A, and triple-negative breast cancer cell line, MDA-MB-231, were obtained from our cell depository. MDA-MB-231 cells were cultured in Dulbecco’s modified Eagle’s medium (DMEM) supplemented with 10% fetal bovine serum (Gibco, Grand Island, NY, USA) and antibiotics (penicillin and streptomycin) (Gibco, Grand Island, NY, USA). MCF-10A cells were cultured in DMEM/F-12 supplemented with GlutaMAX-1, 20 ng/mL recombinant human EGF (Thermo Fischer), 0.5 mg/mL hydrocortisone (Sigma), 100 ng/mL cholera toxin (Sigma), 10 µg/mL insulin (Sigma), 5% horse Serum (Invitrogen) and antibiotics (penicillin and streptomycin) (Gibco, Grand Island, NY, USA).

### 5.7 Morphological Assessment of Breast Cell Response to Mechanical Stimulation

Cell morphology and cluster formation were analysed after 4 days of culture. For analysis, cells were fixed in 4% paraformaldehyde for 10 minutes at room temperature and rinsed 3 times with PBS, waiting 5 minutes for each wash. Afterwards, cells were permeabilized with 0.1% Triton X-100 for 10 minutes at room temperature and rinsed 3 times with PBS, waiting 5 minutes for each wash. For analysis of cell morphology, cells were stained with Alexa Fluor 488 Phalloidin (Invitrogen, Germany) overnight at 4°C and rinsed 3 times with PBS, waiting 5 minutes for each wash. Cells were imaged using a Leica Stellaris SP8 equipped with a 10x objective (N.A = 0.40). The images were 1024x1024 pixels, and z-stacks were taken of 150 μm-thick sections. Cell morphology was visualized using Alexa Fluor 488 Phalloidin signal. Aspect ratio (major axis/minor axis) and roundness (4*area/(π major axis^2)) was determined using ImageJ free hand tool and shape descriptors. Cluster formation was manually counted, whereby cells without contact with other cells were counted as single cells. For morphological and cluster formation analysis, at least 9 stacks were taken per sample, and 3 independent experiments were analysed per condition. Representative higher-resolution images were captured with a 40X (N.A = 0.95) oil immersion objective.

### 5.8 Statistics

Statistical analysis was performed using Prism 10.0 GraphPad software. The significance level was set to p < 0.05 for all conditions. The number of independent biological replicates, sample sizes analyzed, and statistical tests used are stated in the figure legends.

## Author Contributions

Data curation, T.V.; Formal analysis, T.V.; Investigation, Q.W and H.S.Z.; Resources, Q.W and H.S.Z.; Supervision, Q.W and H.S.Z.; Writing—original draft, H.S.Z., Q.W., and T.V.; Writing— review and editing, H.S.Z., Q.W., and T.V. All authors have read and agreed to the published version of the manuscript.

## Acknowledgements

The authors would like to thank Dr. Catherine Whittington and Athenia Jones for gifting MDA-MB-231 cell line and Dr. Michele Vitolo for gifting the MCF-10A cell line.

## Funding Source

This research did not receive any specific grant from funding agencies in the public, commercial, or non-profit sectors.

## Conflicts of Interest

The authors declare no conflicts of interest.

## Data Availability Statement

All data supporting the findings of this study are available from author with reasonable request.

## Notes

### Competing Interest Statement

The authors have declared no competing interest.

### Summary of Updates

The manuscript has been revised to more accurately describe the relationship between collagen matrix stiffness and pore size. The terminology of complete independent control has been moderated to describe substantial decoupling and independent modulation within a defined range. The Discussion has also been expanded to acknowledge that polymerization temperature may influence additional matrix properties, including fibril thickness, collagen organization, network connectivity, and local mechanical properties. The limitations of bulk storage modulus measurements have been further discussed, including the potential contributions of local and time dependent mechanical properties experienced by embedded cells. The Discussion now also acknowledges that cellular remodeling during the four day culture period may alter pore size, fiber organization, and matrix mechanics. Higher quality confocal images have been added to Figure 1. A new Figure 2 has been added to illustrate the image processing and segmentation workflow used for quantitative pore size analysis, including representative raw, processed, binary, and three dimensional Euclidean Distance Transform images. The Methods section has been expanded to describe the image processing procedure. The Introduction and Discussion have been revised to further explain the significance of tuning stiffness and pore size using pure collagen I without chemical crosslinkers or additional matrix components. Proposed mechanisms underlying cellular responses have been reframed as possible mechanisms rather than direct conclusions because they were not directly measured. The statistical analysis has been revised to address potential pseudoreplication. Independent biological gels are now defined as the experimental unit, with individual cell measurements treated as nested within biological gels. Cell morphology data were reanalyzed using nested one way ANOVA with Sidak multiple comparisons. Figure legends were revised to distinguish biological replicates from individual cell measurements, and Figures 2 and 3 now present SD rather than SEM. The Conclusion has been expanded to include future directions. Additional methodological details have been added, including MCF10A culture conditions and the neutralization medium. Grammatical, typographical, and stylistic errors have also been corrected throughout the manuscript.

