## Supplemental File for "Decoupling Bulk Stiffness and Pore Size in 3D Rat Tail Collagen I Matrices"

Theadora Vessell$a^{1}$, Qi We$n^{2}$, Hong Susan Zho$u^{1}$

1 Department of Chemical Engineering, Worcester Polytechnic Institute, 100 Institute Rd, Worcester Massachusetts 01609, USA

2 Department of Physics, Worcester Polytechnic Institute, 100 Institute Rd, Worcester Massachusetts, 01609, USA

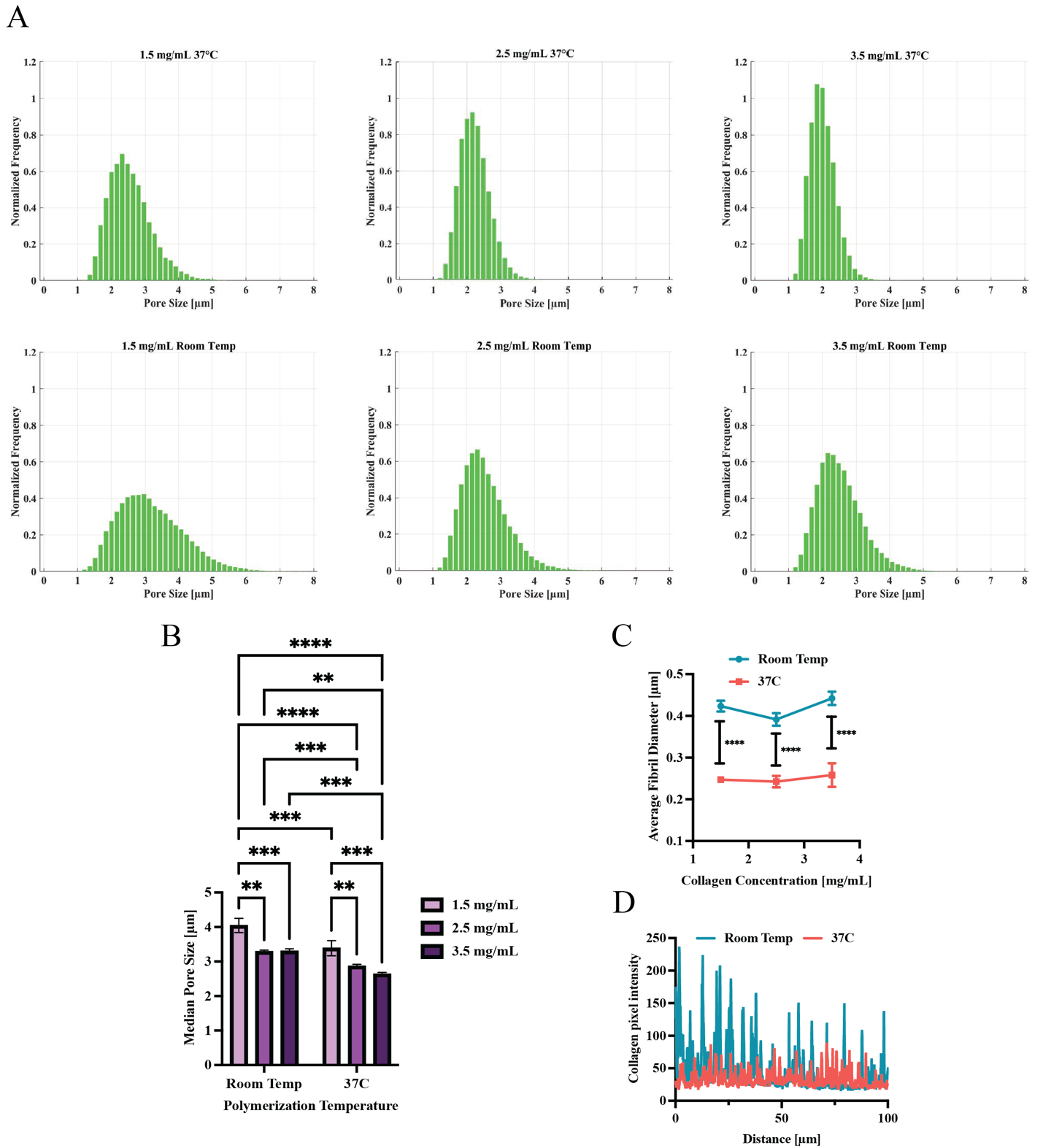

**Supplemental Figure 1. Room temperature collagen polymerization increases median pore size in 3D collagen hydrogels.** (A) Representative average histogram distributions of pore size diameters from 3D pore size analysis.

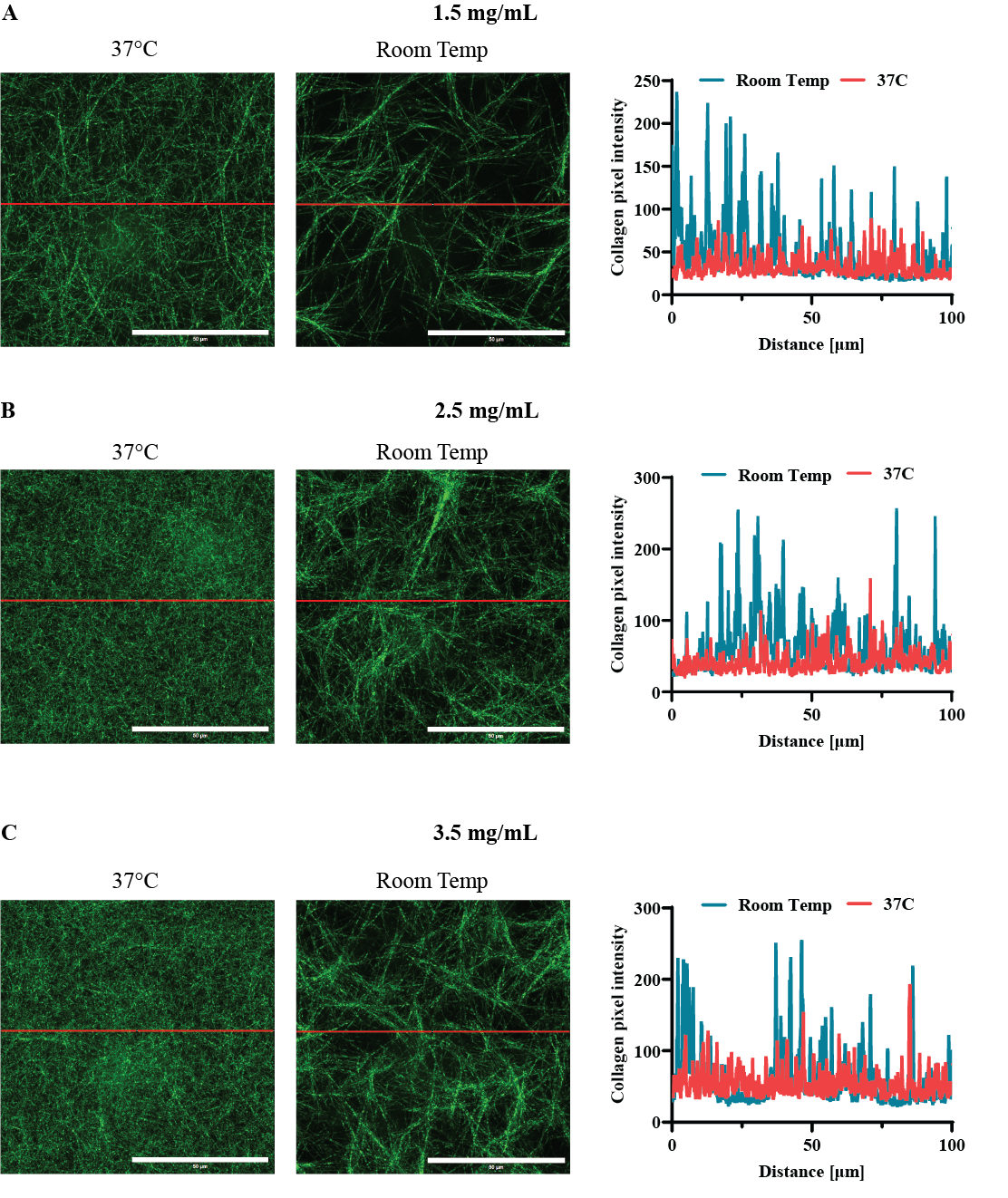

**Supplemental Figure 2. Line scans of mean fiber diameter for all collagen concentrations.** Line scans (representative line in red) for 37°C and Room Temp polymerization temperature for (A) 1.5 mg/mL, (B) 2.5 mg/mL and 3.5 mg/mL Coll I matrices. Scale bar represents 50 µm.

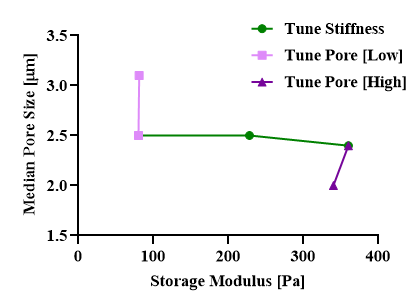

**Supplemental Figure 3. Independent tuning of 3D Coll I matrix stiffness and pore size.** Median pore size [μm] plotted as a function of storage modulus [Pa] for matrices with varied collagen concentration and polymerization temperature. Green circles indicate the condition where stiffness is tuned independently from pore size. Light purple squares represent low stiffness with tuned pore size. Dark purple triangles represent higher stiffness with tuned pore size.

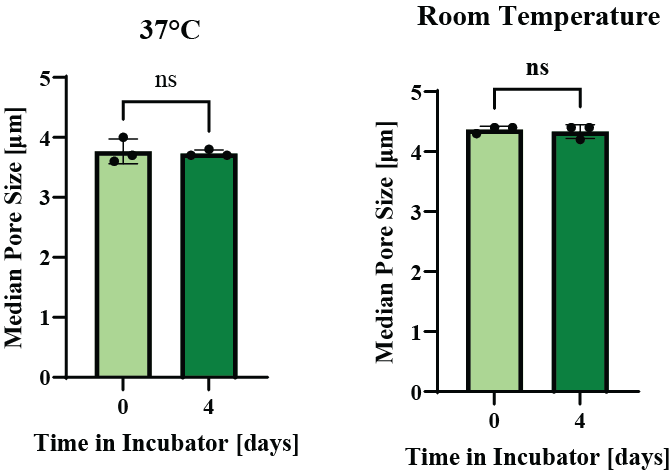

**Supplemental Figure 4. 1.5 mg/mL** **Coll I architecture over time.** Comparison of average median pore diameters on day 0 and day 4. Un-paired t-test, not significant (ns) *P* > 0.05, **P* ≤ 0.05, ***P* ≤ 0.01, ****P* ≤ 0.001, *****P* ≤ 0.0001. Error bars represent SD.

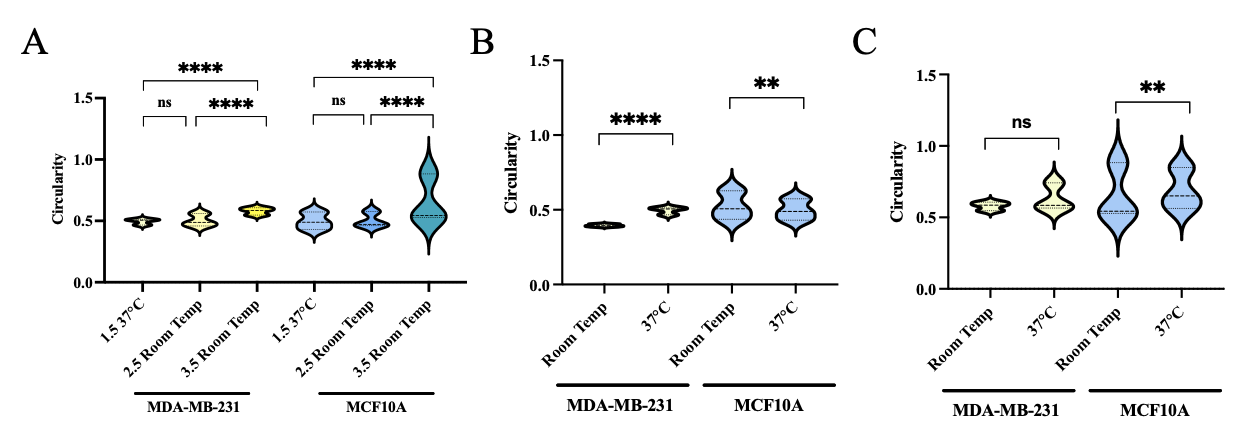

**Supplemental Figure 5. Impact of mechanical stimulation on cell circularity.** Quantitative analysis of MDA-MB-231 breast cancer and MCF-10A breast epithelial cell circularity (1 = perfect circle) as a function of A) stiffness and pore size at both B) low (1.5 mg/mL) and C) high (3.5 mg/mL) concentrations. Significance was tested between samples of varying stiffness and similar pore size. **N** = 3 independent gels per condition; MDA-MB-231 n = 455–1125 cells, MCF-10A n = 387–687 cells. Nested one-way ANOVA followed by Sidak’s multiple-comparisons test was used, with individual measurements nested within biological gels. Not significant (ns) p > 0.05, p ≤ 0.05, *p ≤ 0.01, **p ≤ 0.001, ***p ≤ 0.0001.
